# High-Content Screening to Identify Inhibitors of Dengue Virus Replication

**DOI:** 10.1101/2023.03.24.534108

**Authors:** Jillian G. Hoffstadt, Jesse W. Wotring, Sam Porter, Benjamin Halligan, Matthew J. O’Meara, Andrew W. Tai, Jonathan Z. Sexton

## Abstract

Dengue Virus (DENV) causes dengue fever, a pandemic-potential disease with currently no FDA-approved antivirals. Additionally, the available vaccine for DENV can increase the risk of severe dengue fever for those who have never had a DENV infection due to antibody-dependent enhancements. Thus, there is an urgent need to identify dengue virus antivirals. Antivirals that target NS4B, the replication compartment forming protein of DENV and the flavivirus family, are a promising new drug class that minimize cytotoxic effects to host cells. Drug-repurposing and high-content screening were leveraged to efficiently identify antivirals likely to inhibit NS4B. Using high-content screening, we quantified the morphological patterns of NS4B and envelope (E) protein expression versus time and developed a viral pseudotime model that was able to predict the infection progression to enable drug screening. We then developed a single cell infection classifier for antiviral efficacy and performed high-throughput drug screening of 960 compounds. We identified four concentration-dependent inhibitors of DENV with nanomolar potencies including: Nexium, Pralatrexate, GW4064, and LY411575. LY411575, a gamma secretase inhibitor, exhibited an IC_50_ of 72nM and reduced percent infection to levels indistinguishable from the mock infection control.

## Introduction

The global risk of dengue fever has grown dramatically in recent years, with an estimated 100-400 million annual infections and 96 million clinically severe cases.^1^ Mild cases are flu-like, but without early detection and access to proper medical care, including managing fever with acetaminophen and maintaining body fluid volume, it can progress to dengue shock syndrome (DSS) or dengue hemorrhagic fever (DHF).^2^ DHF symptoms include severe bleeding, organ failure and/or plasma leakage, which can lead to death.^2^ In some regions of Asia and Latin America, severe dengue is a leading cause of serious illness and death.^1^

Dengue fever is caused by dengue virus (DENV) infection, a positive-strand RNA virus in the *Flavivirus* genus. DENV has four different serotypes. If one is infected by one serotype and recovers, they develop lifelong immunity to that serotype of DENV, but due to antibody-dependent enhancements across the four serotypes, subsequent re-infections of another serotype increase the risk of developing severe dengue.^3^ Vaccination can also increase the risk of severe dengue if infected post-immunization.^3^ The DENV vaccine, CYD-TDV, is available in endemic areas and approved in Europe (2018) and the US (2019), but is only recommended for individuals who have DENV seropositivity.^3^

Therefore, there is an urgent need to identify antiviral drugs effective against DENV. Currently, there are no FDA-approved antiviral therapies available for any of the four serotypes of DENV. **Table 1** shows clinical candidates that have reached clinical trials for DENV, including active and failed trials. There were two small molecules in phase 1 clinical trials found to be ineffective^4,5^ and three drugs in phase 2, the most notable being JNJ-A07, a pan serotype dengue virus inhibitor targeting NS3-NS4B viral protein interaction.^6–8,^ Many traditional antiviral drug classes cause toxicity due to off-target effects, whereas antivirals that target viral proteins not found in host cells, like NS4B, are an exciting new therapeutic strategy. Thus, there is an urgent need to diversify anti-dengue candidates, and phenotypic screening is an approach that provides a rapid mechanism of action elucidation.

**Table 1.**
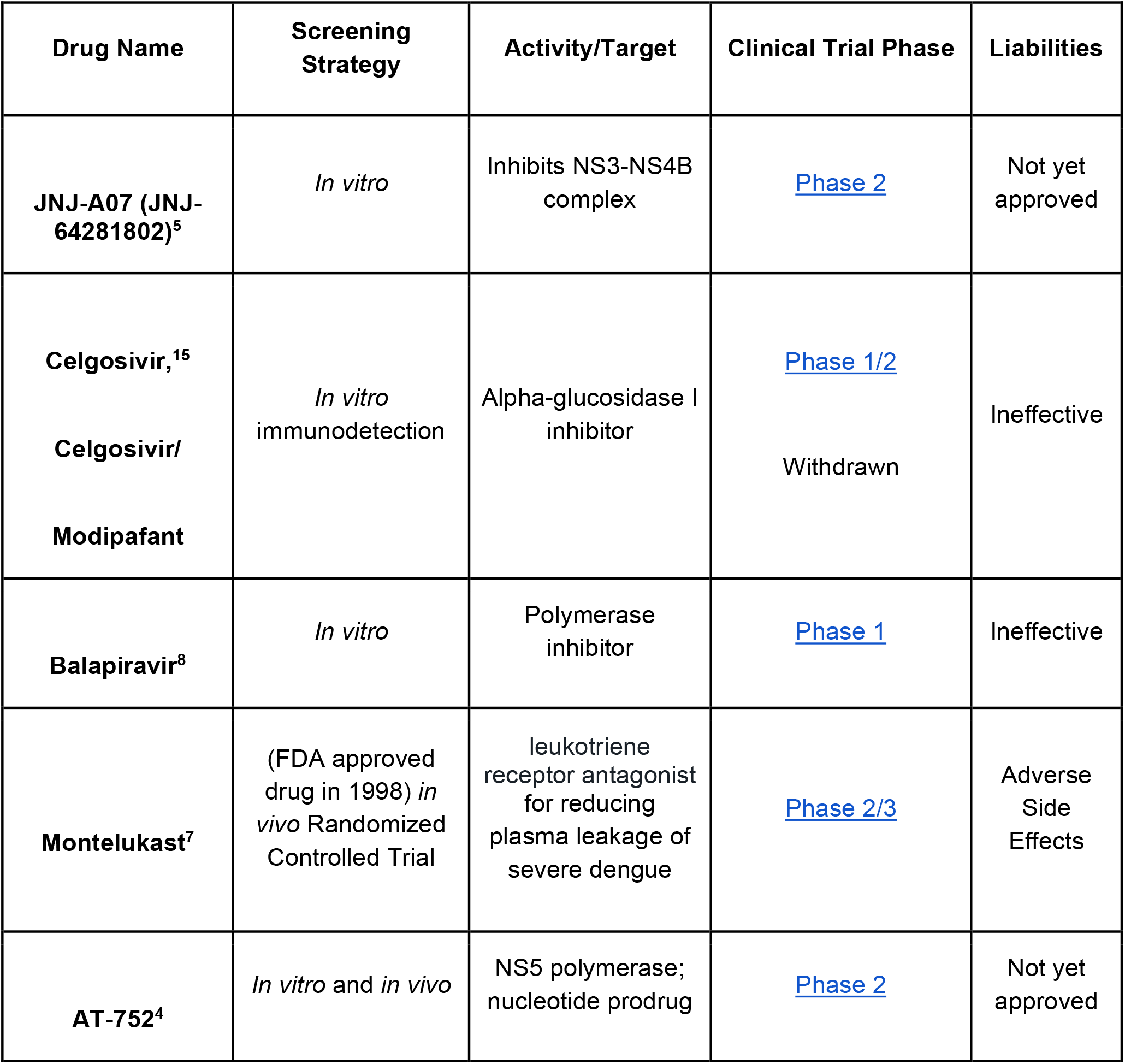
Clinically evaluated compounds for DENV. This table represents a list of all small molecules currently in or were in a clinical trial for DENV. Despite the list of compounds in clinical trials, there is still a need for more chemical diversity of potential DENV inhibitors due to the nature of drug discovery that 80-90% of all drugs will fail to pass clinical trials and become a widespread therapeutic.^17^ Additionally, chemical diversity can identify novel mechanisms and help with challenges due to resistance.

| Drug Name | Screening Strategy | Activity/Target | Clinical Trial Phase | Liabilities |
| --- | --- | --- | --- | --- |
| **JNJ-A07 (JNJ-64281802)<sup>5</sup>** | *In vitro* | Inhibits NS3-NS4B complex | Phase 2 | Not yet approved |
| **Celgosivir,<sup>15</sup>**<br>**Celgosivir/**<br>**Modipafant** | *In vitro*<br>immunodetection | Alpha-glucosidase I inhibitor | Phase 1/2<br>Withdrawn | Ineffective |
| **Balapiravir<sup>8</sup>** | *In vitro* | Polymerase inhibitor | Phase 1 | Ineffective |
| **Montelukast<sup>7</sup>** | (FDA approved drug in 1998) *in vivo* Randomized Controlled Trial | leukotriene receptor antagonist for reducing plasma leakage of severe dengue | Phase 2/3 | Adverse Side Effects |
| **AT-752<sup>4</sup>** | *In vitro* and *in vivo* | NS5 polymerase; nucleotide prodrug | Phase 2 | Not yet approved |

Phenotypic screening is a promising methodology that has been fruitful in identifying small molecule inhibitors. High-content imaging is an extension of phenotypic screening that provides vast amounts of quantitative data about every cell in an experiment by utilizing automated multi-fluorescent confocal microscopy, whereby images are analyzed using a high-throughput computational workflow.^9^ Conventional screening assays are limited in that they only focus on a small number of hand-picked endpoints, whereas morphological profiling allows scientists to draw conclusions from hundreds to thousands of cellular features.^9^ Thus, high-content imaging presents a unique opportunity to better understand the life cycle of DENV replication and how to stop replication. To screen for DENV antivirals more precisely, we first aimed to understand the morphology of DENV replication by visualizing NS4B as well as the envelope (E) protein over 48 hours. Then, through high-content screening, we aimed to identify inhibitors of DENV and provide justification for their mechanism of action.

This phenotypic screening approach has delivered several promising leads for DENV in the past.^7,10^ For example, a phenotypic anti-dengue screen with an eGFP readout conducted by a team of scientists from KU Leuven in partnership with Janssen Pharmaceutica identified JNJ-A07 as a potent inhibitor of NS3-NS4B protein interaction in VeroE6 cells.^7^ Interestingly, this was a broad phenotypic screen and they validated NS4B as a target for DENV by chance. JNJ-A07 is currently in clinical trials for DENV infection.^7^ Our study uses high-content fluorescence imaging, which employs a more targeted approach to discover inhibitors of NS4B by identifying morphological endpoints indicating the disruption of the replication complex.

A promising approach for identifying new therapies with the potential for rapid deployment is drug repurposing, whereby compounds with already established safety profiles are used to treat diseases other than their approved indications. In this study, we explore repurposing drugs as potential clinical therapeutics for DENV. Additionally, many of the repurposed drugs used in this screen are orally bioavailable, accomplishing our goal of finding accessible therapeutics for low-resource countries affected by DENV. Drug repurposing coupled with phenotypic endpoints and high-throughput screening can reveal mechanisms of action for hit compounds because we can compare and contrast phenotypes of drugs with known targets.

Overall, we hypothesize that by using drug repurposing and high-content fluorescence imaging, we will identify DENV replication compartment inhibitors targeting the NS4B protein. Additionally, we aim to describe morphological features of the DENV replication life cycle in the absence of drug treatments to better understand the role of NS4B and the structural envelope protein (E) during replication.

## Methods

### Compounds

The University of Michigan Drug Repurposing Library (DRL) was assembled in the Sexton laboratory from Selleckchem and Cayman Chemical and prepared as 2 mM stock solutions in dimethylsulfoxide (DMSO) in Echo PP Plates. 960 compounds from the DRL were dispensed using Biotek Echo Liquid Handler. Nexium, Pralatrexate, SB271046, IMD0354, LY411575, GW788388, GW4064, and Z-FA-FMK were plated in triplicate dose-response from 8 uM–2nM in DMSO via Biotek Echo Liquid Handler, which makes titrations to find IC_50_ values.

### Cells and Virus

The human hepatoma-derived Huh7.5.1 cells were maintained in Dulbecco’s Modified Eagle Medium (DMEM) supplemented with 10% fetal bovine serum (FBS) and 1X penicillin-streptomycin solution (15140122, Gibco), and were grown at 37 °C with 5% CO_2_ following standard cell culture procedures. To generate wild-type DENV-2 virus, the pD2/IC-30P-NBX plasmid was linearized with XbaI restriction enzyme digestion, in vitro transcribed, and capped with the m7G(5′)ppp(5′)A cap analog using T7 Megascript. This RNA was transfected into Vero E6 cells using TransIT mRNA reagent stocks were grown in Vero E6 cells and titers were determined by TCID_50_ using the Reed and Muench method. All the work with live DENV virus was performed in biosafety level 2 laboratories (BSL2) with the approval of the University of Michigan’s Department of Environment and Health and Safety and the Institutional Biosafety Committee.

### Anti-DENV High Content Bioassays

For drug screening, compounds were dispensed into empty 384-well plates (6057300, Perkin-Elmer, Waltham, MA, USA) using a Beckman Coulter Echo 650 acoustic liquid handler. All wells were normalized to a constant DMSO concentration of 0.2%, and plates contained both infected and uninfected control wells. Huh7.5.1 cells were seeded on top of the compounds at a density of 3,000 cells per well in 50 μL of media. After 24 hours of cell attachment and compound pre-incubation at 37°C and 5% CO_2_, cells were inoculated with DENV at a multiplicity of infection (MOI) of 2. Cells were incubated with virus and compounds for an additional 48 h and then fixed with 100% methanol for 10 mins at -20 °C. Cells were then permeabilized with 0.3% Triton-X100 for 15 minutes. The plates were then incubated with an anti-NS4B primary antibody (GTX103349, GeneTex, Irvine, California, USA) at a dilution of 1:2000 overnight at +4 °C. The staining buffer was PBS with 1.5% w/v BSA, 1% w/v goat serum, and 1X TBS Tween 20. Following primary antibody staining, cells were then stained with 1:1000 secondary antibody Alexa647 for NS4B labeling (A-31573, donkey anti-rabbit, Thermo Fisher, Waltham, MA, USA), 1:4000 CellMask Orange (H32713, Thermo Fisher, Waltham, MA, USA) for cytoplasmic labeling, 1:25 Concanavalin A (C11252, Thermo Fisher, Waltham, MA, USA) for ER labeling, and 1:1000 Hoechst-33342 (H1399, Thermo Fisher, Waltham, MA, USA) for nuclear labeling.

For time series analysis, the plates were treated as described above and were stained with anti-NS4B protein primary antibody at a dilution of 1:2000 and a custom anti-4G2 E protein primary antibody produced in-house^11^ at a dilution of 1:10 overnight at +4 °C. Following primary antibody staining, cells were stained with 1:1000 secondary antibody Alexa647, 1:4000 CellMask Orange, 1:1000 secondary antibody Alexa488 for E protein labeling (A-21121, goat anti-mouse, Thermo Fisher, Waltham, MA, USA), and 1:1000 Hoechst-33342. All dyes were incubated for a total of 45 mins at room temperature. Cells were washed and stored in Phosphate-buffer saline (PBS) before imaging.

### High Content Imaging

Stained assay plates were imaged using a Yokogawa Cell Voyager 8000 (CV8000) microscope with 20X/1.0NA water immersion lens. Maximum intensity projection images were collected over a 10 μm Z-height with 2 μm spacing. A total of 9 fields per well were imaged for all assay plates, accounting for roughly 80% of the total well area. Laser power and camera exposure times were adjusted to yield optimal signal to noise.

### Image Processing

Image processing was performed using the CellPose 2.0 segmentation software (diameter=40, no edge, model=cyto 2) and CellProfiler 4.0 (CellProfiler C. McQuin et al., CellProfiler 3.0: Next-generation image processing for biology. PLoS Biol. 16, e2005970 (2018)). Pipelines were used to identify nuclei (Hoechst), whole cell (HCS CellMask Orange), and measure viral protein intensity and texture features (Alexa Fluor 647 & 488) as well as general cell morphologic measurements. 342 features (Texture, Intensities, Cell Shape, and Area) were collected for each cell. Approximately 100 infected and 100 uninfected cells were manually identified using the CellProfiler Analyst 3.0.4 and used to train a Random Forest model to classify all cells as infected/uninfected in each plate. The model performed with high accuracy (>99.8%) under 5-fold cross-validation.

### Time-Series Analysis

Cell-level features of 0 hours post-infection (HPI) and 48 HPI for the NS4B and E protein (“NS4B & E pro” model) image channels were used to train an XGBoost decision tree model to score infection under 5-fold cross-validation performed using Scikit-learn (Scikit-learn: Machine Learning in Python, Pedregosa *et al*., JMLR 12, pp. 2825-2830, 2011.). Only fields with cell counts above 200 were included in the training set. Each time point was replicated in 64 wells, but after filtering for cell count, 44 wells for hour 0 were included, 53 wells for hour 12, 24 wells for hour 20, 22 wells for hour 28, 18 wells for hour 36, and 44 wells for hour 48. To model infection dynamics, separate models for only NS4B (“NS4B only”) and only E protein (“Epro only”) were trained independently. All cells were then scored for infection by each of the three models and averaged to the field level and the image level. Scores of infection were compared for statistical significance in a paired t-test (Kassambara A (2022). rstatix: Pipe-Friendly Framework for Basic Statistical Tests. R package version 0.7.1, <https://CRAN.R-project.org/package=rstatix>).

Cellular features for the NS4B and E protein image channels were also used in a dimensionality reduction analysis via UMAP for X and Y coordinates. Due to the large number of cell observations, a 10% random subset (650,000 cells) was analyzed.

## Results

### Time-Series Analysis of Expression of NS4B and E Protein

Cell morphologic analysis of cells infected with DENV at eight hour increments revealed characteristics of the replication life cycle. We quantified these observations by extracting intensity, texture, and area/shape measurements from viral protein channels in high-content images and analyzed the features with machine learning. For this analysis, the object-level data from the nuclei, cytoplasm, NS4B, and E protein channels were used to produce a UMAP embedding of all cells. The embedding shows distinct clustering of cellular objects based on intensity and texture of E protein. The earlier time points in this experiment captured early replication morphology based on the expression of E protein before NS4B. Cells with little to no expression of viral proteins clustered towards the right-hand side of the graph and cells with high expression of viral proteins clustered towards the left-hand side. The UMAP and representative images of the clusters can be seen in **Figure 1**.

**Figure 1:**
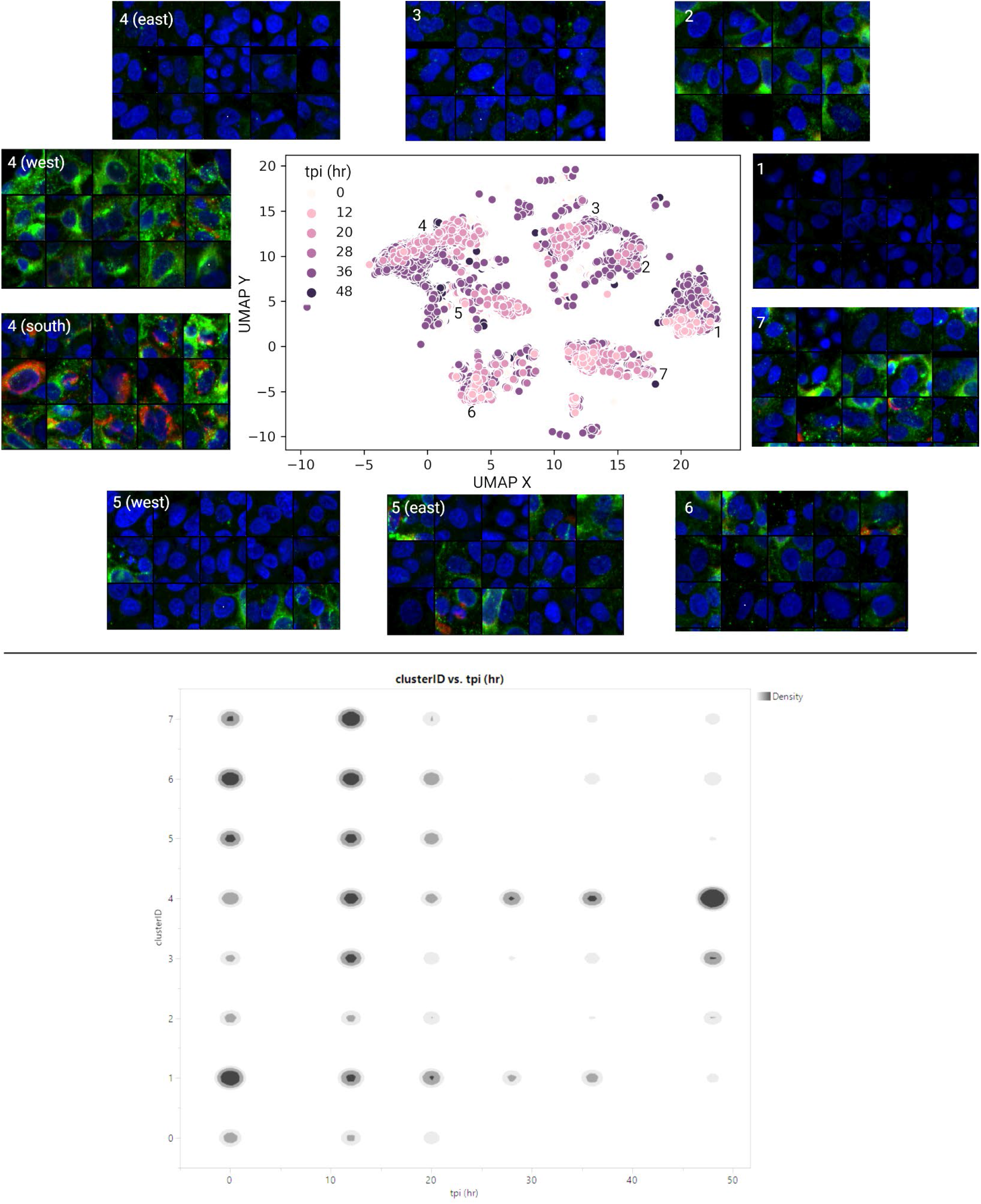
UMAP of cells shows clustering based on time point. The UMAP embedding of cell-level morphological features shows seven distinct clusters of infected cells. Cells from every time point are present in every cluster, but there are strong preferences for certain morphology in each cluster. Cluster 1 shows mock infection cells. Cluster 2 shows flat and even E protein texture in the cytoplasm. Cluster 3 shows small speckles of E protein signal in the cytoplasm. Cluster 4 can be divided into an east and west region. The east region of cluster 4 shows heterogeneous texture in the nuclei of uninfected cells. The west region of cluster 4 shows strong intensity for E protein only in infected cells. The southern part of cluster 4 shows strong NS4B and E protein intensity in infected cells. Cluster 5 can again be divided into a west and east region. The west region of cluster 5 shows nuclei close together, which could have resulted in segmentation of multiple nuclei as 1 cell. The east region of cluster 5 shows some low intensity of E protein in most cells. Cluster 6 shows more speckled texture of E protein. Cluster 7 shows a strong E protein intensity coupled with a speckled cytoplasmic texture.

Then, a decision tree XGBoost model was trained on the viral channels of images from hour 0 and 48 as positive and negative controls, respectively. We trained three separate models on both viral channels (NS4B & Epro), NS4B only, and E protein only. We then scored the probability of infection for all cellular objects in every well using each of the three models. The scores were averaged to the image level and finally to the well level. We tested the statistical consistency of staining for NS4B and E protein. From hour 0 to 12 and 0 to 20, the E protein model sees a statistically significant increase in average well-level score, whereas the NS4B model does not. At hour 20, the NS4B model probability score was different from the control model by a statistically significant margin, while the E protein model was not. These results in **Figure 2** suggest that at early points of the replication cycle, DENV expresses E protein before it expresses NS4B protein.

**Figure 2:**
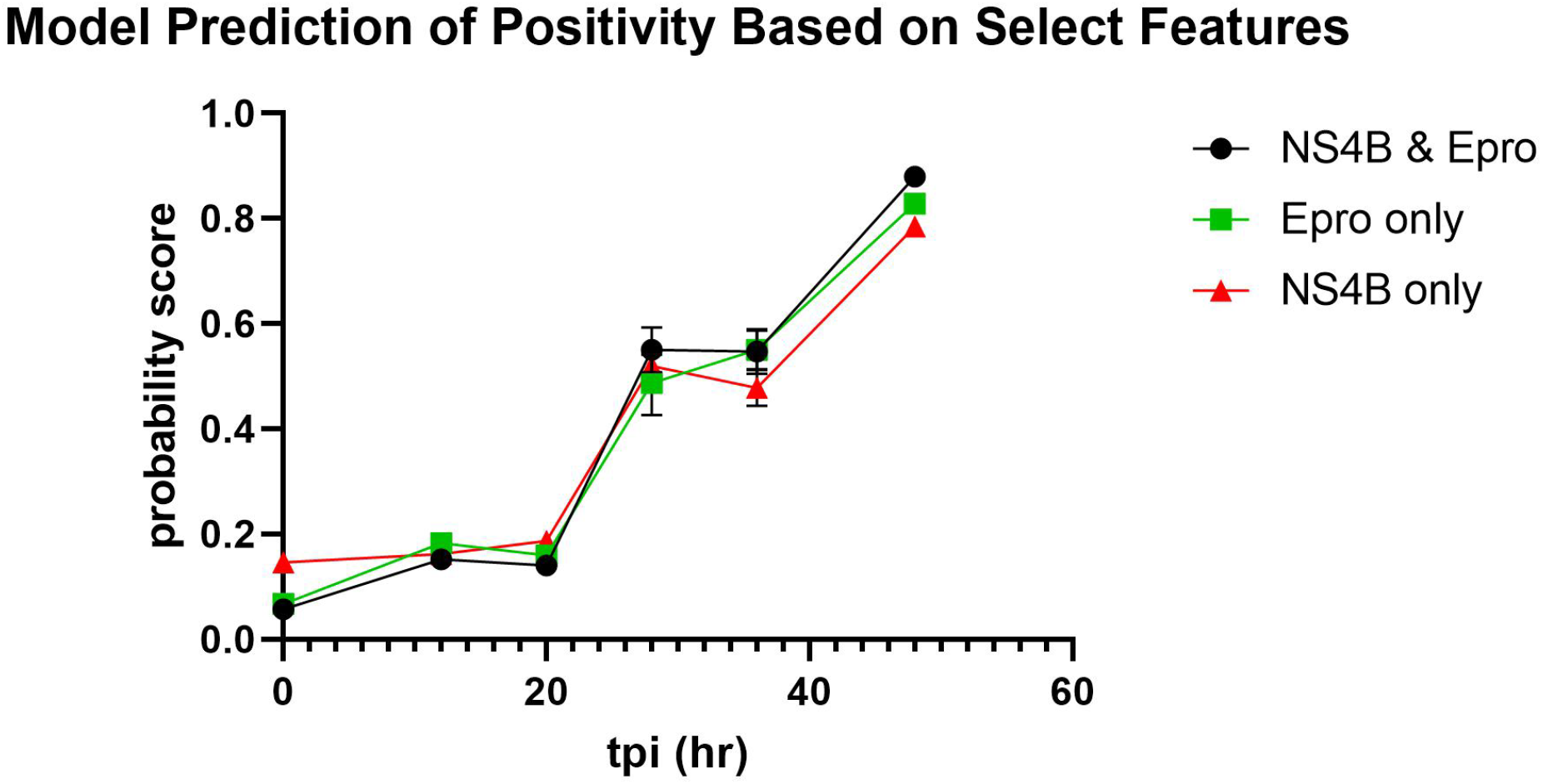
Significant increase in Epro only model probability score at hour 12 shows E protein forms at early stages of DENV lifecycle. The probability score of infection based on select features was averaged to the well level. There was a statistically significant difference in the probability scores of wells from the model trained on E protein only from hour 0 to 12 and 20. There was not a statistically significant difference in the probability scores from the NS4B model for this time interval. All other time intervals, the NS4B model and the E protein model agreed with each other and the control model whether there was or wasn’t a statistically significant change in protein expression. At hour 12, none of the models had a statistically significant difference in their means. At hour 20, the NS4B model probability score was different from the control model by a statistically significant margin, while the E protein model was not.

### High-Content Screening to Identify Replication Compartment Inhibitors

Here, we aimed to identify viral replication compartment inhibitors in a cell model for infection using Huh7.5.1 cells. We designed and optimized a high-content fluorescence imaging assay in 384-well plate format using Huh7.5.1 and measured cell viability and viral inhibition. We screened 960 compounds from the University of Michigan Drug Repurposing Library, including FDA approved drugs and clinical candidates. We used the detection of viral envelope (E) protein and NS4B as markers for DENV infection and cell count per well as an indicator of cell viability. From the primary screen, we selected eight compounds for followup concentration response studies. This assay was highly reproducible and had a Z’ value of 0.7. As summarized in the **Figure 3A** workflow, Huh7.5.1 cells were preincubated with a 10-point 3-fold dilution series of the seven compounds for 24 hours and then infected with wild-type DENV-2 virus and incubated for an additional 48 hours. Following infection, cells were fixed, permeabilized, and stained to identify nuclei, cytoplasm, and viral E and NS4B proteins. Assay plates were imaged at 20X magnification using a Yokogawa CellVoyager 8000 high content imaging platform (*n* = 9 fields captured per well) and processed using the image segmentation software CellPose and image analysis software CellProfiler. CellProfiler Analyst was used to classify cells as positive or negative for viral protein expression using a random forest model to tabulate percent infection per-well. Data from CellProfiler was used to determine percent viability using cell counts per field. Infection data were normalized to the average well-level percent positive for infected controls. Percent viability was determined by normalizing the average well-level cell counts for the infected control.

**Figure 3.**
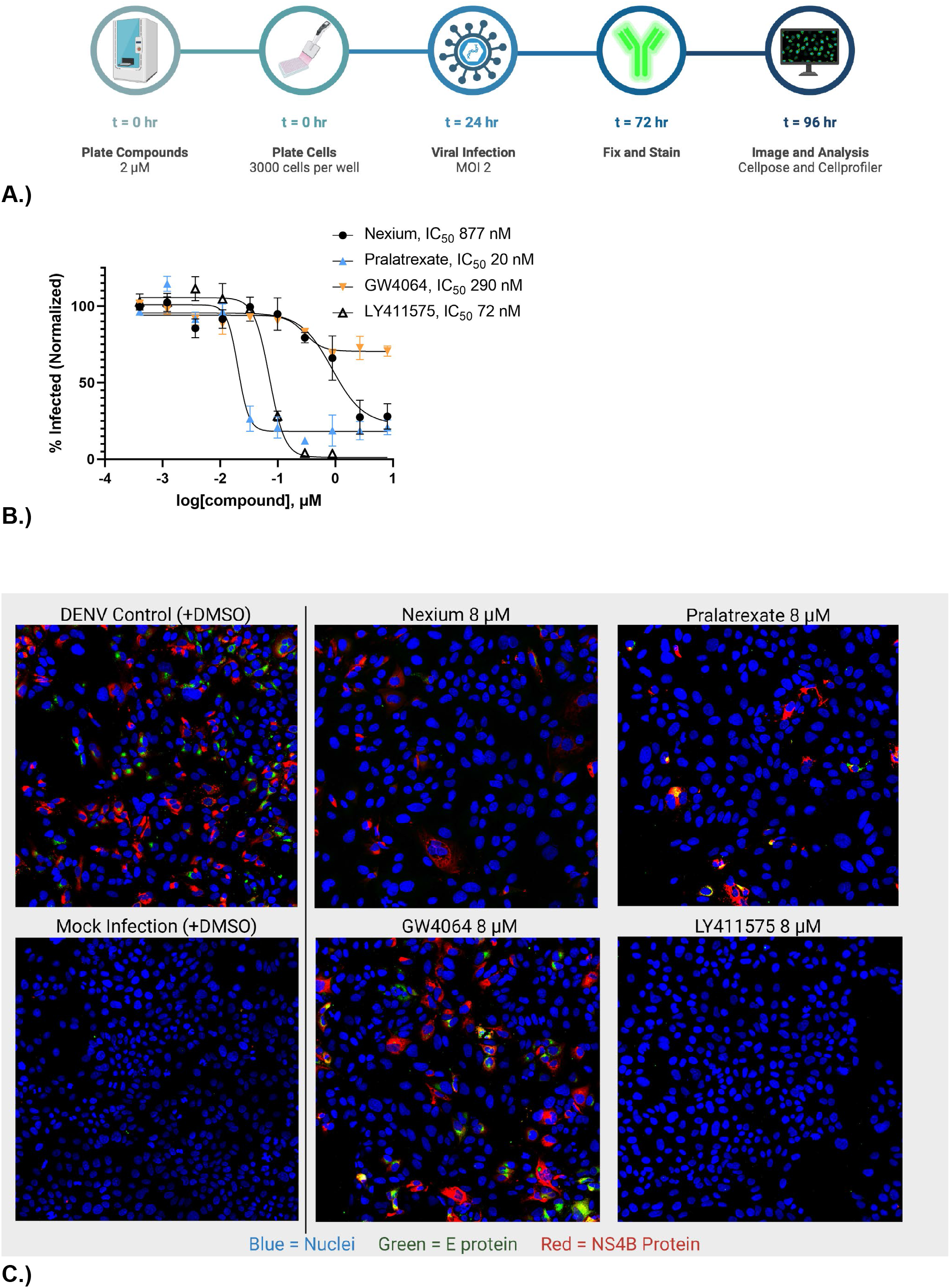
Drug Repurposing Screen for Dengue Antivirals. A.) High-throughput screening workflow. B.) Drug screen dose response. C.) Images of the inhibitors and controls.

We found that four drugs produced dose-dependent inhibition of DENV infection. Nexium, a proton pump inhibitor, had a potent 50% maximal inhibition (IC_50_) value of 877 nM. Pralatrexate, a DHFR inhibitor, had an IC_50_ value of 20 nM. GW4064, a farnesoid X receptor inhibitor, had an IC_50_ value of 290 nM. And, LY411575, a gamma-secretase inhibitor, had an IC_50_ value of 72 nM, bringing the percent positive of infection per well down to almost zero at the highest concentration (8 μM). The concentration-response curves for this experiment are shown in **Figure 3B**. Representative images for infected control, mock, and 8 μM drug conditions are included in **Figure 3C**. In conclusion, we identified four drugs with nanomolar potency to inhibit dengue infection.

## Discussion

There remains an urgent need for DENV therapeutics, which can be used effectively to reduce the burden of severe dengue on developing countries’ health infrastructure. Especially since the current vaccine cannot be supplied to individuals who have never had a DENV infection. Small molecule antivirals could provide a significant benefit to those who may be vaccine hesitant, immunocompromised, or living in areas with limited access to health care.

The formation of E protein before NS4B protein could inform decisions about targets for inhibitor therapeutics. An inhibitor that targets envelope protein formation could be used as an early replication inhibitor given immediately after exposure to DENV. Our results agree with previous findings that report E protein is the first viral protein to make contact with host cells.^12^

We have demonstrated that the FDA-approved drugs Nexium and Pralatrexate and the commercially available GW4064 and LY411575 have potent antiviral activity against DENV infection in human cells. Nexium is a proton-pump inhibitor (PPI) with satisfactory bioavailability used to treat GERD and other conditions causing gastric acid hypersecretion. There has been success to use PPI inhibitors as antivirals for HIV, in which the drug interacts with Tsg101, an ESCRT, and prevents the virus from reaching its budding site on the ER. Other viruses analogously employ Tsg101, including flaviviruses, but Nexium was found to be ineffective in an FFU assay in Vero cells for inhibition of DENV.^13^ Considering the strong concentration-dependent inhibition of DENV by Nexium based on our assay techniques, perhaps the cell line employed by Watanabe et al. was not permissive enough for DENV replication. Pralatrexate is a DHFR inhibitor with satisfactory bioavailability used to treat relapsed or refractory peripheral T-cell lymphoma. In a drug-repurposing screen for SARS-CoV-2, Pralatrexate was found to inhibit SARS-CoV-2, which could mean it is a broad-spectrum antiviral, but the authors did not describe a mechanism of action.^14^ DHFR inhibitors are non-specific to the viral cells versus the host cells,^15^ but there was no significant reduction in cell viability at the concentrations we tested. GW4064 was first described in 2000 as a farnesoid X receptor (FXR) agonist later discovered to have poor bioavailability (10%).^16,17^ FXR is a bile acid receptor. GW4064 was found to inhibit FXR and disrupt lipid homeostasis in rotaviruses, which the authors propose could be a valid mechanism to target the DENV capsid protein because it is made of lipids.^16^ Due to GW4064’s poor bioavailability, other FXR agonists would be great candidates for a new drug class of antivirals. LY411575 is an agonist of the catalytic domain of gamma-secretase, presenilin, and has been shown to inhibit HCV, which is a flavivirus family member of DENV.^18^ LY411575 arrests HCV replication by inhibiting a presenilin-like aspartic protease, signal peptide peptidase (SPP), from cleaving the C-terminal transmembrane region of the core protein during maturation.^18^ This is compelling because DENV also recruits SPP to cleave its structural proteins between the capsid and prM.^19^ There is no experimental data available for the oral bioavailability of LY411575, however it has demonstrated in vivo bioactivity.^20^

This study has important limitations. As a whole, high-content imaging is subject to noise and background signals that can cause overfitting of machine learning models and clustering to an unimportant source of noise. Luckily, there are methods to remove background signal by subtracting a percent of total signal from every image via CellProfiler. Additionally, by only labeling two viral proteins as our readout, we are limited in our ability to draw strong conclusions about the mechanism of action, so target elucidation of our hits is necessary. Although cell-based assays provide better transition to *in vivo* than biochemical assays, there is still a risk that the lead compounds could show no bioactivity *in vivo*. Of our compounds, Pralatrexate is a rather aggressive chemotherapeutic agent that may not be wise to prescribe to very ill patients unless the dose can be tolerated. And, GW4064 has such low bioavailability that this compound would not be a good clinical candidate. Finally, further testing is needed to confirm if our hits are acting on-target.

Nonetheless, this does not distract from the overall contributions of the study. For the first time ever, we reported high-content images of the replication life cycle morphology of cells infected with DENV. The multiple clusters based on E protein texture and intensity could provide insight into the structure and function of E protein. Additionally, morphological profiling can provide insight into the mechanism of action of antivirals, which is a future direction this project should take, perhaps by documenting additional viral proteins in a more robust cell-painting assay. Ultimately, we described four potent potential antivirals for DENV and confirmed two novel mechanisms of action to arrest DENV replication: FXR and gamma-secretase agonists.

## Conclusion

There is still an urgent need for effective anti-DENV therapeutics due to limited vaccine efficacy, lack of approved antivirals, and the pandemic-potentiality of dengue fever. We have identified four drugs with clinical translational potential: Nexium, Pralatrexate, LY411575, and GW4064. Nexium and Pralatrexate are already FDA-approved drugs with well-established safety profiles, which would be easy to transition to clinical trials. LY411575 is the most promising of the lead compounds due to its double-digit nanomolar potency and reduction of viral infection to levels indistinguishable from the mock control. We support further target elucidation and *in vivo* testing of LY411575 for dengue virus. Morphological profiling of the replication life cycle visualized the heterogeneous phenotypes of DENV. Our work emphasizes the utility of high-content screening coupled with machine learning approaches to provide strong insight into the efficacy of antiviral drugs.

---

**Supplementary Figure 1.**
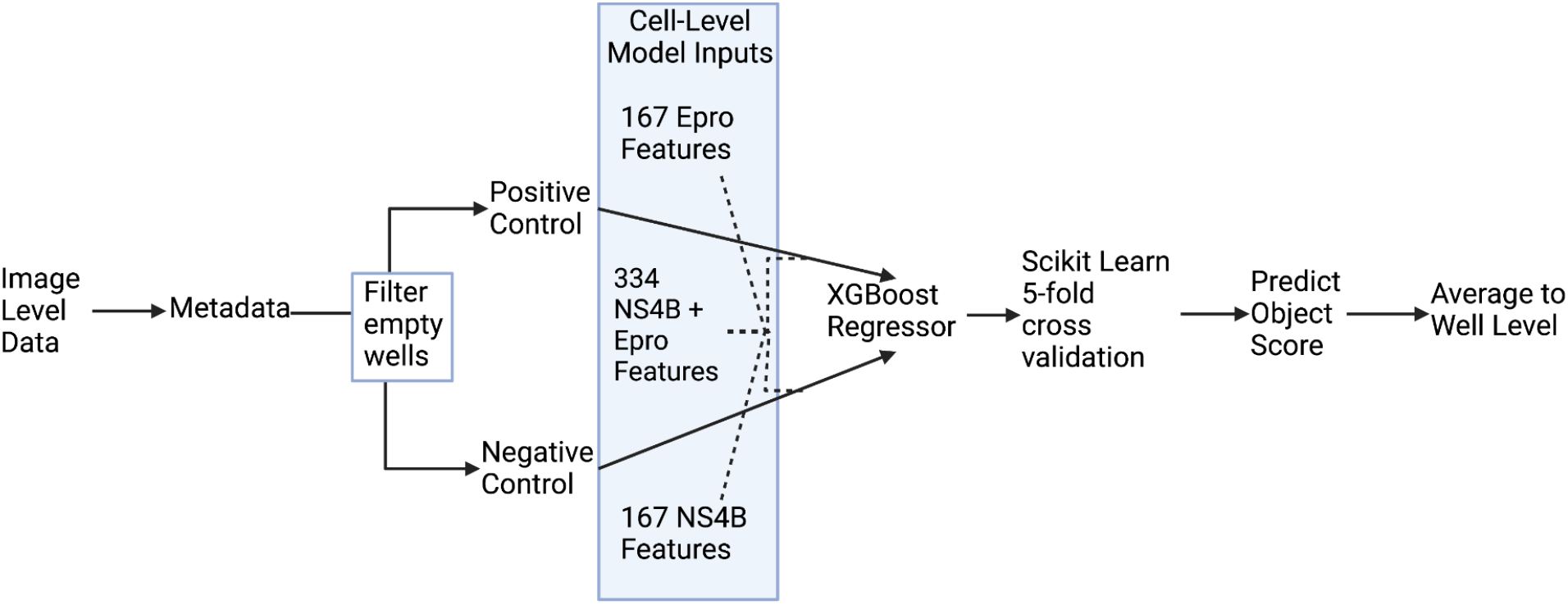
Machine Learning Model Workflow.

## Notes

### Competing Interest Statement

The authors have declared no competing interest.

